# Urothelium-specific expression of mutationally activated *Pik3ca* initiates early lesions of non-invasive bladder cancer

**DOI:** 10.1101/2023.05.22.541489

**Authors:** Lauren Shuman, Jonathan Pham, Thomas Wildermuth, Xue-Ru Wu, Vonn Walter, Joshua I. Warrick, David J. DeGraff

## Abstract

Despite the fact that ∼70% of bladder cancers are non-invasive and have high recurrence rates, early stage disease is understudied. The relative lack of models to validate the contribution of molecular drivers of bladder tumorigenesis is a significant issue. While mutations in *PIK3CA* are frequent in human bladder cancer, an *in vivo* model for understanding their contribution to bladder tumorigenesis is unavailable. Therefore, a *Upk2-Cre/Pik3ca*^*H1047R*^ mouse model expressing one or two *R26-Pik3ca*^*H1047R*^ alleles in a urothelium-specific manner was created. *Pik3ca*^*H1047R*^ functionality was confirmed by quantifying Akt phosphorylation and mice were characterized by assessing urothelial thickness, nuclear atypia, and expression of luminal and basal markers at 6 and 12 months of age. At 6 months, *Pik3ca*^*H1047R*^ mice developed increased urothelial thickness and nuclear atypia, however, at 12 months, *Pik3ca*^*H1047R*^ mice did not exhibit progressive disease. Immunohistochemistry shows urothelium maintained luminal differentiation characterized by high Foxa1 and Pparγ expression. In addition, mice were subjected to low-dose carcinogen exposure (*N*-Butyl-*N*-(4-hydroxybutyl)nitrosamine (BBN)). Surprisingly, *Pik3ca*^*H1047R*^ mice exhibited no significant differences after exposure relative to mice without exposure. Furthermore, ssGSEA analysis of invasive human tumors showed those with mutant *PIK3CA* do not exhibit significantly increased PI3K/AKT pathway activity compared to wildtype *PIK3CA* tumors. Overall, these data suggest that *Pik3ca*^*H1047R*^ can elicit early tumorigenic changes in the urothelium, but progression to invasion may require additional genetic alterations.

## Introduction

An estimated ∼570,000 new cases of bladder cancer (BC) are diagnosed worldwide every year, with about 82,000 new cases and 17,000 deaths occurring in the United States.^1, 2^ Clinically, BC is classified as non-muscle invasive or muscle invasive disease and this distinction is useful in determining if aggressive treatment is warranted.^3^ However, classifying BC as non-invasive or invasive can help to distinguish the unique biological characteristics of each as distinct entities and therefore identify appropriate therapies.^3-5^ Approximately 70% of newly diagnosed BC cases are non-invasive BC which include papillary tumors (stage Ta) and carcinoma *in situ* (CIS).^4, 6^ In addition to comprising the majority of diagnoses, non-invasive BC also has a high recurrence rate (up to 70%) which requires long term monitoring and treatment.^7^ The high recurrence rate therefore is a major contributing factor to the high cost of BC treatment, making BC one of the most costly cancers to manage.^8^

Despite the challenges of clinically managing non-invasive BC, most research is focused on invasive disease. One reason for this is that the availability of models for the study of early stage disease is limited. Establishment of early stage models of BC is vital to furthering our understanding of events leading to tumor initiation and progression, and will offer *in vivo* platforms for the development of novel treatments, while additionally broadening the scope of BC research to increase focus on non-invasive disease. Currently, transgenic and conditional/tissue-specific knockout mouse models provide platforms to test the contribution of genetic alterations (e.g., somatic mutation and DNA copy number loss) to tumor development and progression. However, additional clinically relevant models of early stage BC are required.

The phosphatidylinositol-4,5-bisphosphate 3-kinase catalytic subunit alpha (*PIK3CA*) gene encodes the p110α catalytic subunit of phosphoinositide-3-kinase (PI3K). Signal transduction following activation of the PI3K/AKT pathway plays an important role in normal cell biology and pathologic states.^9, 10^ While *PTEN* deletion is common in invasive BC and cooperates with other common genetic alterations in advanced disease,^11-14^ mutations in *PIK3CA* are more frequently detected and have been identified across all BC stages. In fact, mutations have been found in 20-30% of BC independent of stage, indicating these mutations are an early event.^11, 15-17^ While many activating mutations in *PIK3CA* have been detected in BC, an *in vivo* model of activating *Pik3ca* mutations in BC does not exist. Of the most common activating mutations, H1047R is currently available to study in a commercial mouse model. Therefore, we utilized this strain for the creation of a novel mouse model of BC driven by *Pik3ca* mutation.

## Materials and Methods

### Mouse breeding, genotyping, and experiments

All animal experiments were performed in accordance with and following approval of the Institutional Animal Care and Use Committee (IACUC) at Pennsylvania State University College of Medicine. To generate mice with urothelium-specific expression of one or two mouse *Pik3ca*^*H1047R*^ mutant alleles, commercially available *R26-Pik3ca*^*H1047R*^ mice (stock 016977; The Jackson Laboratory; Bar Harbor, ME) were bred with previously described *Upk2-Cre* mice which express Cre recombinase in a urothelium-specific manner.^18, 19^ This breeding strategy resulted in the development of male and female mice with one (*Upk2-Cre/Pik3ca*^*H1047R+/-*^) or two (*Upk2-Cre/Pik3ca*^*H1047R+/+*^) alleles of mouse mutant *Pik3ca* plus genetic control mice (no *Cre* and/or *R26-Pik3ca*^*H1047R*^ alleles).

All mice were genotyped by PCR analysis using the following primers: *Upk2-Cre* Forward - 5’-CGTACTGACGGTGGGAGAAT-3’; *Upk2-Cre* Reverse - 5’-TGCATGATCTCCGGTATTGA-3’; *R26-Pik3ca*^*H1047R*^ Common (oIMR8545, The Jackson Laboratory) - 5’-AAAGTCGCTCTGAGTTGTTAT-3’; *R26-Pik3ca*^*H1047R*^ Mutant Reverse (oIMR8052, The Jackson Laboratory) - GCGAAGAGTTTGTCCTCAACC; *R26-Pik3ca*^*H1047R*^ Reverse (oIMR8546, The Jackson Laboratory) - 5’-GGAGCGGGAGAAATGGATATG-3’. The presence of *Upk2-Cre* was indicated by a single band at ∼350 bp. Wildtype and mutant *R26-Pik3ca*^*H1047R*^ mouse alleles were indicated by bands present at ∼650 bp and 340 bp, respectively.

For experiments, one cohort of mice consisting of genetic control and *Upk2-Cre/Pik3ca*^*H1047R*^ mice with one and two mutant alleles was aged for 6 or 12 months (n = 56) while another cohort consisting of genetic control and *Upk2-Cre/Pik3ca*^*H1047R*^ mice with 2 mutant alleles was aged for 6 months and then exposed to the carcinogen *N*-Butyl-*N*-(4-hydroxybutyl)nitrosamine (BBN; B0938; TCI America; Portland, OR) in their drinking water at a sub-carcinogenic concentration of 0.01% for 10 weeks (n = 25). BBN water was changed twice weekly during the 10-week period. Mice were then euthanized for bladder collection.

### Immunohistochemistry

Dissected bladders were fixed in 10% formalin, processed, and embedded in paraffin using standard procedures. Five micron sections were cut for H&E and immunohistochemical staining. Immunohistochemistry (IHC) was performed as previously described.^20^ Slides were deparaffinized, rehydrated through graded ethanol, then washed in distilled water. Antigen retrieval was performed using 1% antigen unmasking solution (H-3300; Vector Laboratories; Burlingame, CA) and heating slides in a pressure cooker (CPC-600; Cuisinart; Stamford, CT) for 20 minutes at high pressure. Then pressure was released in short bursts to prevent boiling and preserve tissue integrity. Slides were cooled to room temperature (RT) and washed 3 times for 10 minutes in phosphate-buffered saline (PBS; pH 7.4). The subsequent incubations were performed at RT unless otherwise noted. Endogenous peroxidases were blocked using 1% hydrogen peroxide in methanol for 20 minutes and slides were washed again in PBS. Sections were blocked using horse serum (S-2000; Vector Laboratories) diluted in PBS for 1 hour to reduce nonspecific antibody binding and then incubated with primary antibody at 4 °C overnight. Primary antibodies used include rabbit anti-phospho-Akt (S473) (1:200; 4060; Cell Signaling Technology; Danvers, MA), rabbit anti-phospho-Akt (T308) (1:200; 13038; Cell Signaling Technology), goat anti-Foxa1 (1:1000; sc-6553; Santa Cruz Biotechnology; Dallas, TX), rabbit anti-Pparγ (1:200; 2430; Cell Signaling Technology), mouse anti-Krt5/6 (1:200; M7237; Agilent; Santa Clara, CA), and mouse anti-Krt14 (1:200; NCL-L-LL002; Leica Biosystems; Buffalo Grove, IL). Following overnight incubation, slides were washed in PBS and incubated in biotinylated secondary antibody (1:200) diluted in PBS containing horse serum for 1 hour. Secondary antibodies used include anti-rabbit and anti-goat antibodies (BA-1000 and BA-9500; Vector Laboratories). Specific antibody binding was visualized using the Vectastain Elite ABC Peroxidase kit (PK-6100; Vector Laboratories) according to manufacturer’s protocol with diaminobenzidine (DAB; K346811-2; Agilent) as the chromogen. Sections were then counterstained with hematoxylin and mounted with Cytoseal (8310-4; Thermo Fisher Scientific; Waltham, MA). This procedure was modified for mouse primary antibodies for use with the Mouse on Mouse Immunodetection Kit (BMK-2202; Vector Laboratories) according to manufacturer’s protocol. It should also be noted that for phospho-Akt antibodies, Tris-buffered saline with Tween (TBST) and TBS were used for washes and diluent, respectively, in place of PBS throughout the procedure.

### Urothelial thickness and histological scoring

Urothelial thickness was measured using cellSens Entry 1.13 imaging software (Olympus, Center Valley, PA) on H&E sections. Five measurements of the urothelium were taken and averaged per bladder then graphed using GraphPad Prism (Version 9, San Diego, CA). Statistical significance of urothelial thickness for 6- and 12-month aging mouse groups was determined via application of Tukey’s multiple comparisons tests using GraphPad Prism. Statistical significance of urothelial thickness for the comparison of 6-month-old mice with and without BBN exposure was determined by applying two-way ANOVA tests both with and without interaction using R 4.2.0 (The R Foundation, Vienna, Austria).^21^ All tissues were examined by a dedicated genitourinary pathologist (JIW) in a blinded manner. For nuclear atypia, a score of 0-2 was assigned to describe the extent of atypia observed with 0 meaning no atypia, 1 meaning patchy atypia, and 2 meaning extensive atypia seen throughout the urothelium. For phosphorylated Akt (p-Akt) IHC stains, expression levels were scored by calculating the H-score. The H-score was determined by multiplying percentage of positive cells x intensity of staining (0-3). Box plots were generated and Kruskal–Wallis tests were applied to compare the H-scores of p-Akt across mouse groups (controls, one mutant *Pik3ca* allele, and two mutant *Pik3ca* alleles) using R 4.2.0.^21^

### Graphical representation of mutations and ssGSEA analysis of TCGA data

Gene-level RNA sequencing read counts for the TCGA BC cohort (TCGA-BLCA) were downloaded from the Genomic Data Commons.^22^ After restricting to protein-coding genes, the edgeR R package was applied to compute gene-level log counts-per-million reads mapped (CPM) values.^23-25^ With the log CPM values as input, the GSVA R package was used to calculate single sample gene set enrichment analysis (ssGSEA) scores based on the Hallmark PI3K/AKT/MTOR gene set that represent PI3K/AKT/MTOR pathway activity levels in each sample.^26^ For follow-up ssGSEA analysis, missense hotspot mutations recognized as oncogenic within other genes of the PI3K/AKT pathway were identified within TCGA dataset using cBioPortal (www.cbioportal.org, date of last access: 10/25/2022) and samples with these mutations were removed.^27, 28^ Such mutations were identified in the following genes: *FGFR3, ERBB2, ERBB3, HRAS, NRAS, PTEN*, and *AKT1*. Kruskal–Wallis and Wilcoxon rank sum tests were applied to compare the ssGSEA scores across groups of interest (normal tissue, *PIK3CA* wildtype tumors, and *PIK3CA* mutant tumors). A lollipop plot displaying the locations of *PIK3CA* mutations in the TCGA BC cohort was created using the maftools R package and gene mutation information in the combined multicenter MAF.^29, 30^ These analyses were performed using R 4.2.0.^21^

## Results

### *Upk2-Cre/Pik3ca*^*H1047R*^ mice express increased levels of nuclear p-Akt

The development of a novel mouse model of early urothelial tumorigenesis was initiated by crossing a previously described *Upk2-Cre* mouse line with a commercially available *R26-Pik3ca*^*H1047R*^ mouse line.^18, 19^ This resulted in a new strain of *Upk2-Cre/Pik3ca*^*H1047R*^ mice which constitutively express one or two alleles of *Pik3ca*^*H1047R*^ in a urothelium-specific manner (Figure 1A). *Cre* is expressed in the luminal and intermediate cells allowing for the stop codon (flanked by *loxp* sites) preceding the *Pik3ca* mutant gene to be excised, resulting in expression of *Pik3ca*^*H1047R*^.

**Figure 1.**
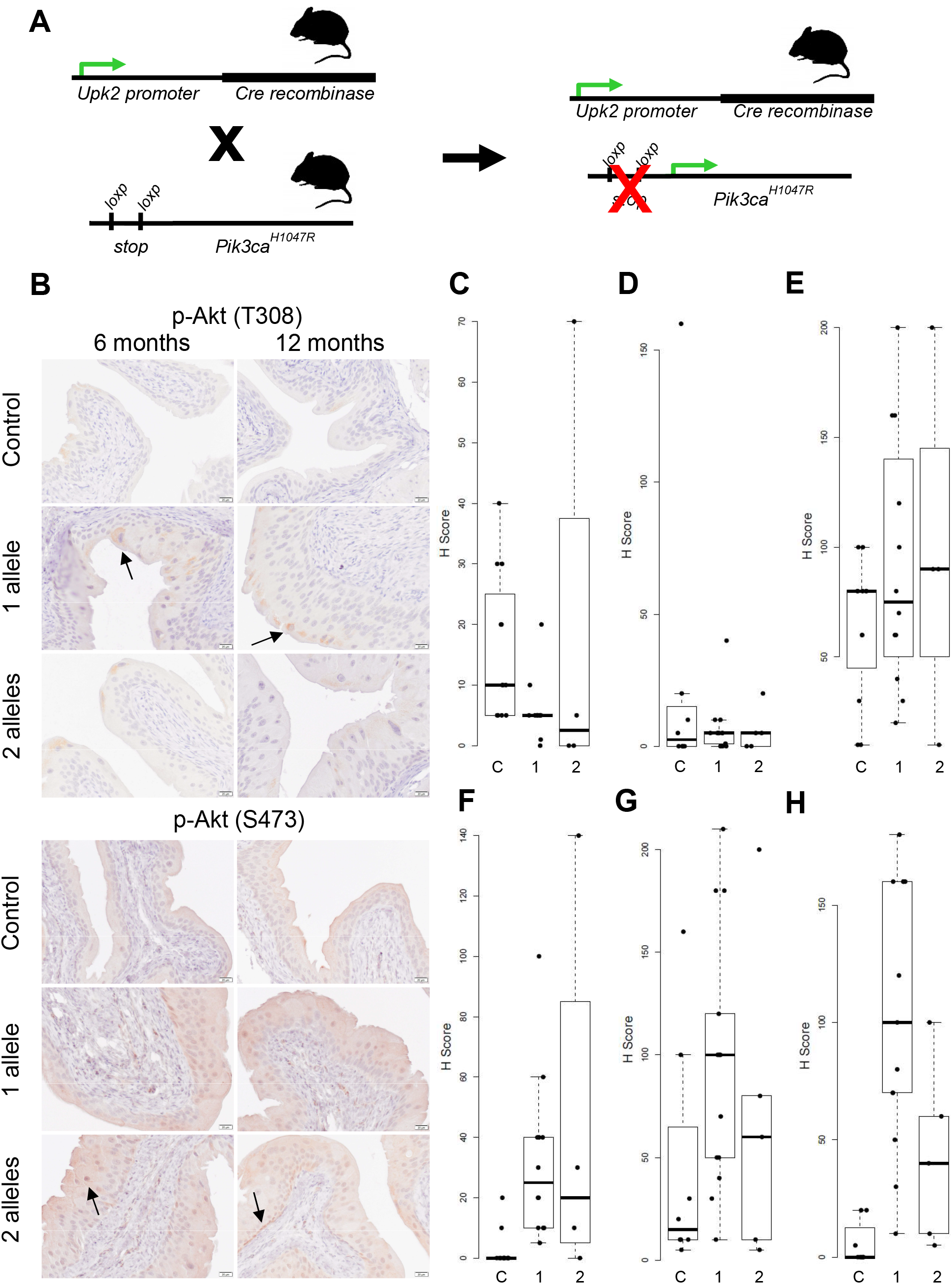
*Pik3ca*^*H1047R*^ functionality is confirmed by increased phosphorylation of Akt in mutant mice. (A) A schematic detailing the breeding strategy used in the creation of *Upk2-Cre/Pik3ca*^*H1047R*^ mice. Previously described *Upk2-Cre* mice were crossed with commercially available *R26-Pik3ca*^*H1047R*^ mice to obtain a novel *Upk2-Cre/Pik3ca*^*H1047R*^ mouse model expressing one or two alleles of *R26-Pik3ca*^*H1047R*^ specifically in the urothelium. (B) Immunohistochemical staining of p-Akt (T308 and S473) in *Upk2-Cre/Pik3ca*^*H1047R*^ mouse urothelium. Examples of cytoplasmic T308 staining of luminal cells as well as nuclear S473 staining are indicated by arrows. Scale bar = 20 µm. (C-H) Quantification of p-Akt (T308 and S473) IHC staining of *Upk2-Cre/Pik3ca*^*H1047R*^ mice. For the 6-month-old age group: control n = 12, 1 allele n = 12, and 2 allele n = 4. For the 12-month-old age group: control n = 8, 1 allele n = 13, and 2 allele n = 5. (C) Cytoplasmic T308 staining of 6-month-old mice (p = 0.0359; Kruskal–Wallis). (D) Cytoplasmic T308 staining of 12-month-old mice (p = 0.9350; Kruskal–Wallis). (E) Cytoplasmic S473 staining of 6-month-old mice (p = 0.506; Kruskal–Wallis). (F) Nuclear S473 staining of 6-month-old mice (p = 0.000526; Kruskal– Wallis). (G) Cytoplasmic S473 staining of 12-month-old mice (p = 0.0989; Kruskal–Wallis). (H) Nuclear S473 staining of 12-month-old mice (p = 0.000339; Kruskal–Wallis). Annotations: C - control, 1 - 1 allele, 2 - 2 alleles.

The functionality of *Pik3ca*^*H1047R*^ expression was confirmed by assessing PI3K/AKT pathway activation. Phosphorylated levels of Akt (p-Akt) were determined by scoring IHC staining of amino acid residues T308 and S473 (Figure 1B-H). Both cytoplasmic and nuclear staining were assessed. Most notably, nuclear staining of S473 was significantly increased in the mutant mice compared to controls at 6 and 12 months of age (p = 5.26e-4 and p = 3.39e-4, respectively; Kruskal–Wallis). While significant increases in cytoplasmic T308 or S473 staining were not observed, it is interesting that cytoplasmic T308 staining was observed in umbrella and intermediate cells in which the *Upk2* promoter driving *Cre* is active (examples shown with arrows). Nuclear staining of T308 was not observed. Overall, this data confirms that expression of *Pik3ca*^*H1047R*^ activates downstream signaling *in vivo*.

### Expression of *Pik3ca*^*H1047R*^ in urothelial cells results in hyperplasia and nuclear atypia

Morphologic analysis showed 6- and 12-month-old mice with one or two alleles of *Pik3ca*^*H1047R*^ developed urothelial hyperplasia, as indicated by increased urothelial thickness (Figure 2A). To quantify the extent of hyperplasia in the *Upk2-Cre/Pik3ca*^*H1047R*^ mice, urothelial thickness (µm) was measured at 6 and 12 months of age (Figure 2B). At 6 months, urothelial thickness was significantly increased in mice with one and two alleles of *Pik3ca*^*H1047R*^ compared to controls (p-value = 1.0e-4 and < 1.0e-4, respectively; Tukey’s multiple comparisons). At 12 months of age, urothelial thickness was also significantly increased in mice with one and two alleles of *Pik3ca*^*H1047R*^ compared to controls (p-value < 1.0e-4 and = 5.0e-4, respectively; Tukey’s multiple comparisons). Urothelial thickness was progressive, with mutant mice developing thicker urothelium at 12 months of age.

**Figure 2.**
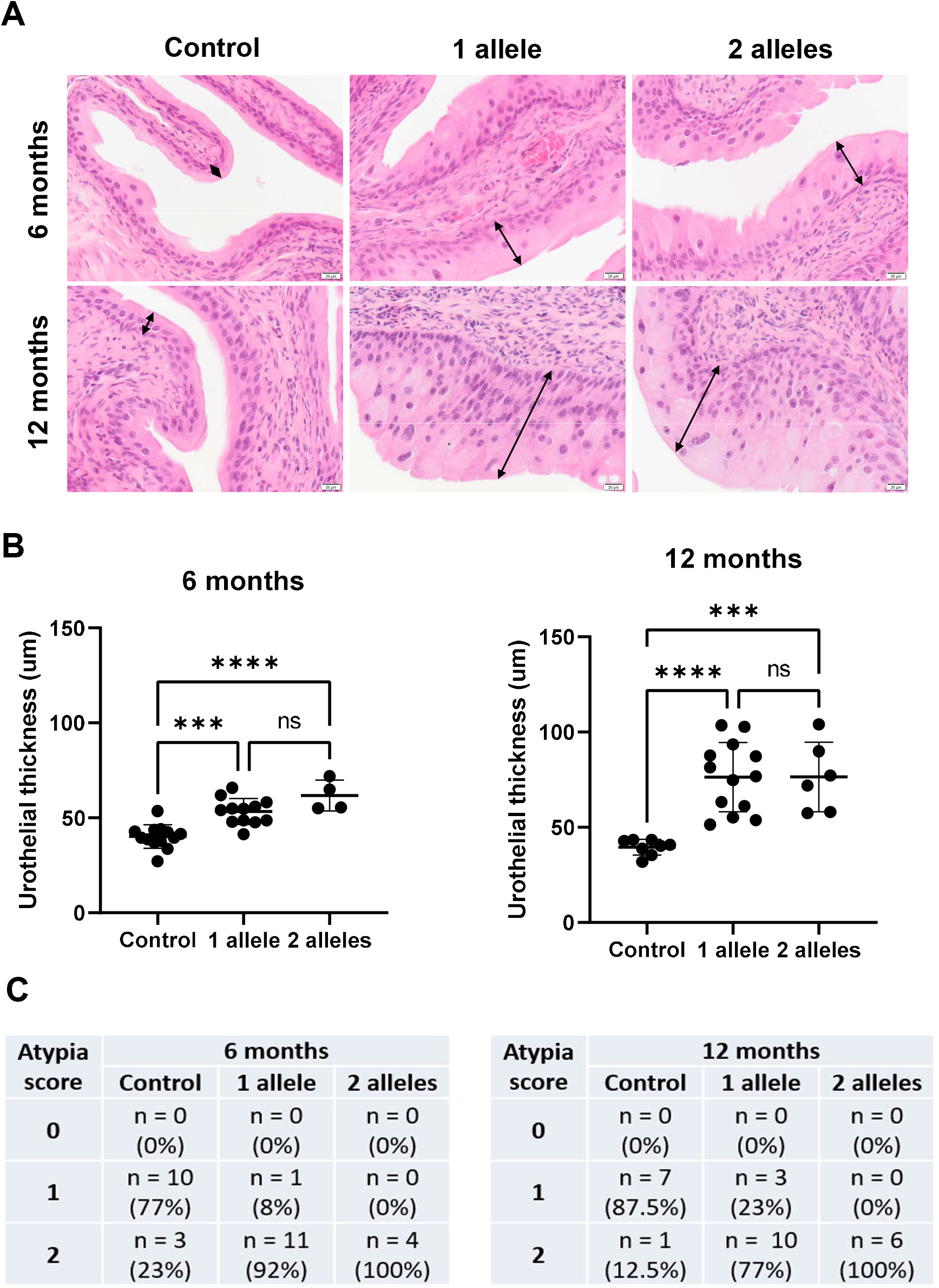
*Upk2-Cre/Pik3ca*^H1047R^ mice exhibit increased urothelial thickness and nuclear atypia. (A) Representative H&E images of control and *Upk2-Cre/Pik3ca*^*H1047R*^ mice with one or two alleles of mutant *Pik3ca* aged for 6 and 12 months. *Upk2-Cre/Pik3ca*^*H1047R*^ mice develop thicker urothelium after 6 and 12 months of aging. The thickness of the urothelium is indicated using arrows. Scale bar = 20 µm. (B) Dot plots illustrating urothelial thickness (µm) of *Upk2-Cre/Pik3ca*^*H1047R*^ mice at 6 and 12 months of age. At 6 months, urothelial thickness is significantly increased in mice with one (n = 12) and two (n = 4) alleles of *Pik3ca*^*H1047R*^ compared to controls (n = 13) (p-value = 0.0001 and < 0.0001, respectively; Tukey’s multiple comparisons). Urothelial thickness is also significantly increased in mice aged 12 months with one (n = 13) and two (n = 6) alleles of *Pik3ca*^*H1047R*^ compared to control mice (n = 8) (p-value < 0.0001 and = 0.0005, respectively; Tukey’s multiple comparisons). Error bars show mean with standard deviation. Annotations: *** - p-value ≤ 0.001, **** - p-value ≤ 0.0001, ns - not significant. (C) Tables summarizing the nuclear atypia scores of *Upk2-Cre/Pik3ca*^*H1047R*^ mice at 6 and 12 months of age. At 6 months, atypia is increased in mice with one (n = 12) and two (n = 4) alleles of *Pik3ca*^*H1047R*^ compared to control (n = 13). Atypia is also increased in mice aged 12 months with one (n = 13) and two (n = 6) alleles of *Pik3ca*^*H1047R*^ compared to control mice (n = 8).

Nuclear atypia was also observed in *Upk2-Cre/Pik3ca*^*H1047R*^ mice (Figure 2C). To quantify nuclear atypia, a score of 0-2 was assigned to describe the extent of atypia observed (0 = no atypia; 1 = patchy atypia; and 2 = extensive atypia seen throughout the urothelium). At 6 months of age, atypia was clearly increased in mice with one and two alleles of *Pik3ca*^*H1047R*^ compared to controls. Atypia was also increased in mice at 12 months of age with one and two alleles of *Pik3ca*^*H1047R*^ compared to controls. Notably, atypia was observed largely within the luminal and intermediate layers of the urothelium which were targeted by the *Upk2* promoter driving *Cre*, coinciding with p-Akt described in Figure 1.

### The urothelium of *Upk2-Cre/Pik3ca*^*H1047R*^ mice exhibits high expression of luminal differentiation markers

To determine whether the *Upk2-Cre/Pik3ca*^*H1047R*^ mouse model exhibited a luminal or basal profile, a set of luminal (Foxa1 and Pparγ) and basal (Krt5/6 and Krt14) differentiation markers was utilized for IHC analysis (Figure 3). Overall, mice with one and two alleles of *Pik3ca*^*H1047R*^ maintained a high level of Foxa1 and Pparγ expression similar to urothelium of genetic controls at 6 and 12 months of age. Also, like genetic control urothelium, mutant mice expressed low levels of Krt5/6 and Krt14. These observations suggest *Upk2-Cre/Pik3ca*^*H1047R*^ mice retained a luminal expression profile similar to that observed in human early stage BC.

**Figure 3.**
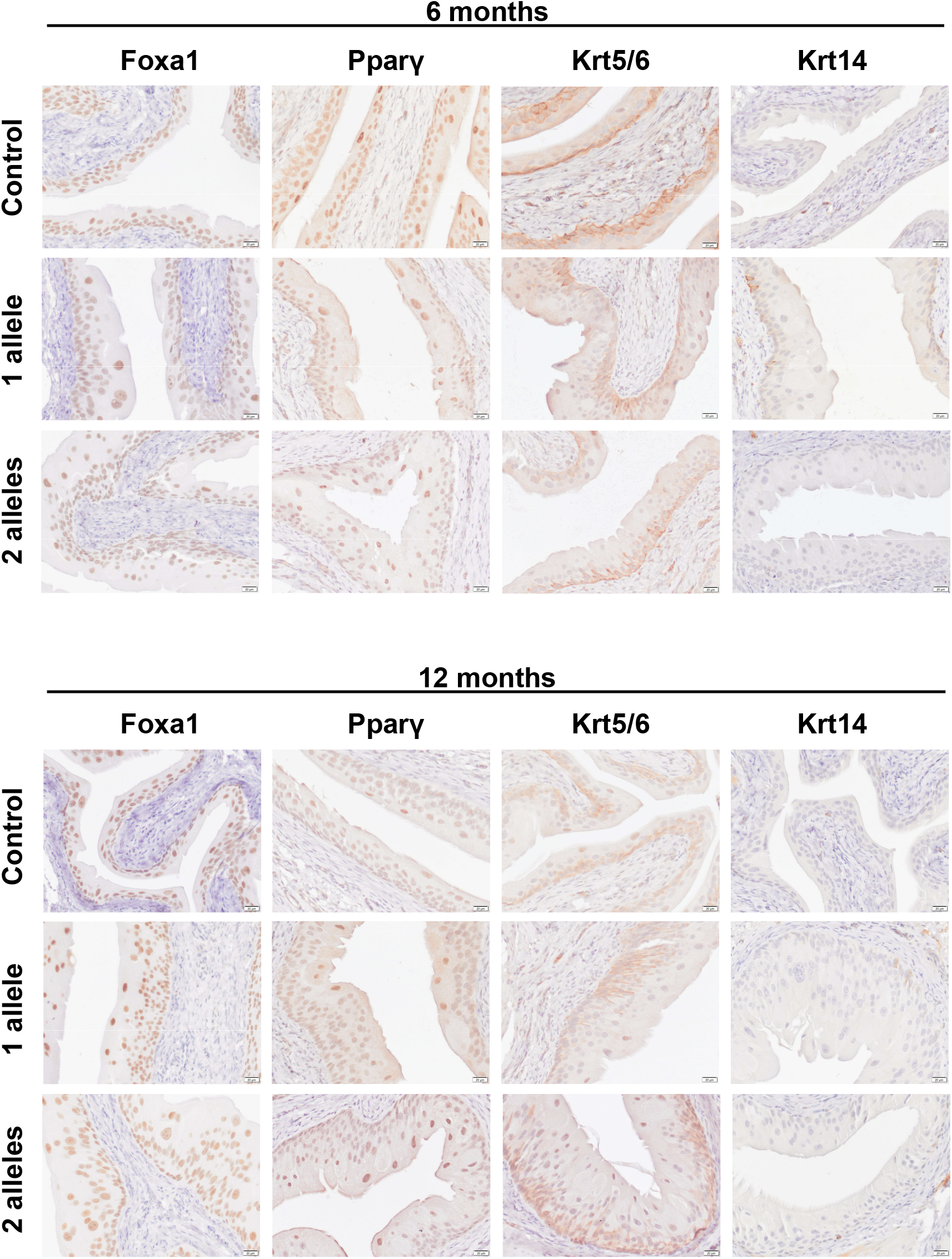
Luminal differentiation markers Foxa1 and Pparγ are highly expressed in *Upk2-Cre/Pik3ca*^H1047R^ mouse urothelium. Immunohistochemical staining of luminal (Foxa1 and Pparγ) and basal (Krt5/6 and Krt14) markers of urothelial differentiation in *Upk2-Cre/Pik3ca*^*H1047R*^ mice aged 6 and 12 months. High expression of luminal markers Foxa1 and Pparγ is maintained in the urothelium of mice with one and two alleles of *Pik3ca*^*H1047R*^. In addition, basal markers Krt5/6 and Krt14 remain low in expression. Scale bar = 20 µm.

### *Upk2-Cre/Pik3ca*^*H1047R*^ mice do not exhibit enhanced susceptibility to the carcinogen BBN

In order to determine the impact of *Pik3ca*^*H1047R*^ expression on sensitivity to a bladder-specific carcinogen, 6-month-old control and *Upk2-Cre/Pik3ca*^*H1047R+/+*^ mutant mice were exposed to a sub-carcinogenic concentration of BBN (0.01% via drinking water for 10 weeks). Mutant mice exposed to BBN showed thicker urothelium compared to control mice exposed to BBN (Figure 4A). In order to separately assess the effects of *Pik3ca*^*H1047R*^ mutations and BBN treatment, quantitative measurements of urothelial thickness were plotted for control and mutant mice, both with and without BBN treatment (Figure 4B). A two-way ANOVA model with interaction was fit using factors for *Pik3ca*^*H1047R*^ mutations and BBN treatment. The interaction term was not significant (p = 0.3941), suggesting that the effect of BBN treatment is similar in control and mutant mice. The two-way ANOVA model was then re-fit without the interaction term. The marginal test of equality for BBN was not statistically significant (p = 0.2097), suggesting that BBN treatment did not alter urothelial thickness after controlling for the effect of *Pik3ca*^*H1047R*^ mutations. However, the corresponding marginal test for *Pik3ca*^*H1047R*^ mutations was highly significant (p = 2.024e-6). Clear differences in nuclear atypia were also not detected (Figure 4C). Additionally, IHC staining for Foxa1, Pparγ, Krt5/6, and Krt14 was again conducted and showed mutant mice maintain similar patterns of the luminal and basal markers compared to controls following exposure to BBN (Figure 5). These results indicate *Upk2-Cre/Pik3ca*^*H1047R+/+*^ mice did not exhibit increased sensitivity to a sub-carcinogenic concentration of BBN.

**Figure 4.**
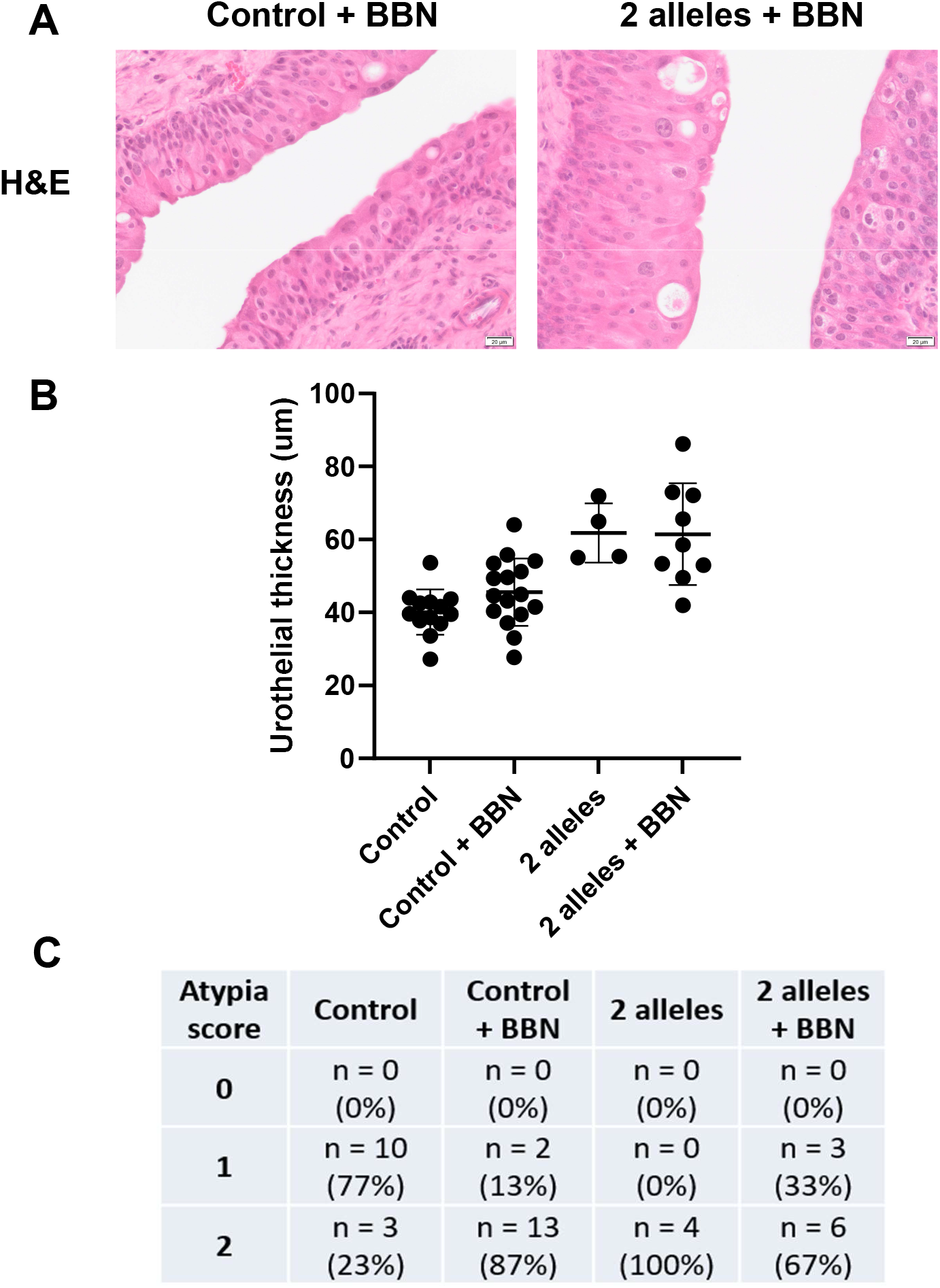
*Upk2-Cre/Pik3ca*^H1047R^ mice do not show significant differences in susceptibility to BBN. Six-month-old control and mutant mice with two alleles of *Pik3ca*^*H1047R*^ were given 0.01% BBN in their drinking water for 10 weeks. These mice were then compared to the 6-month-old mice previously assessed which were not exposed to BBN. (A) H&E staining shows slightly thicker urothelium of mutant mice with similar morphology to controls after exposure to BBN. Scale bar = 20 µm. (B) Urothelial thickness (µm) measurements show no significant difference driven by *Pik3ca* mutation in combination with BBN exposure (p = 0.3941; two-way ANOVA with interaction). The two-way ANOVA model was then re-fit without the interaction term. The marginal test for BBN is not significant (p = 0.2097), however, the marginal test for *Pik3ca*^*H1047R*^ mutations is highly significant (p = 2.024e-6). The numbers of mice per group are as follows: control n = 13, control + BBN n = 16, 2 alleles n = 4, and 2 alleles + BBN n = 9. Error bars show mean with standard deviation. (C) Clear differences in nuclear atypia are not observed following exposure to BBN. The numbers of mice per group are as follows: control n = 13, control + BBN n = 15, 2 alleles n = 4, and 2 alleles + BBN n = 9.

**Figure 5.**
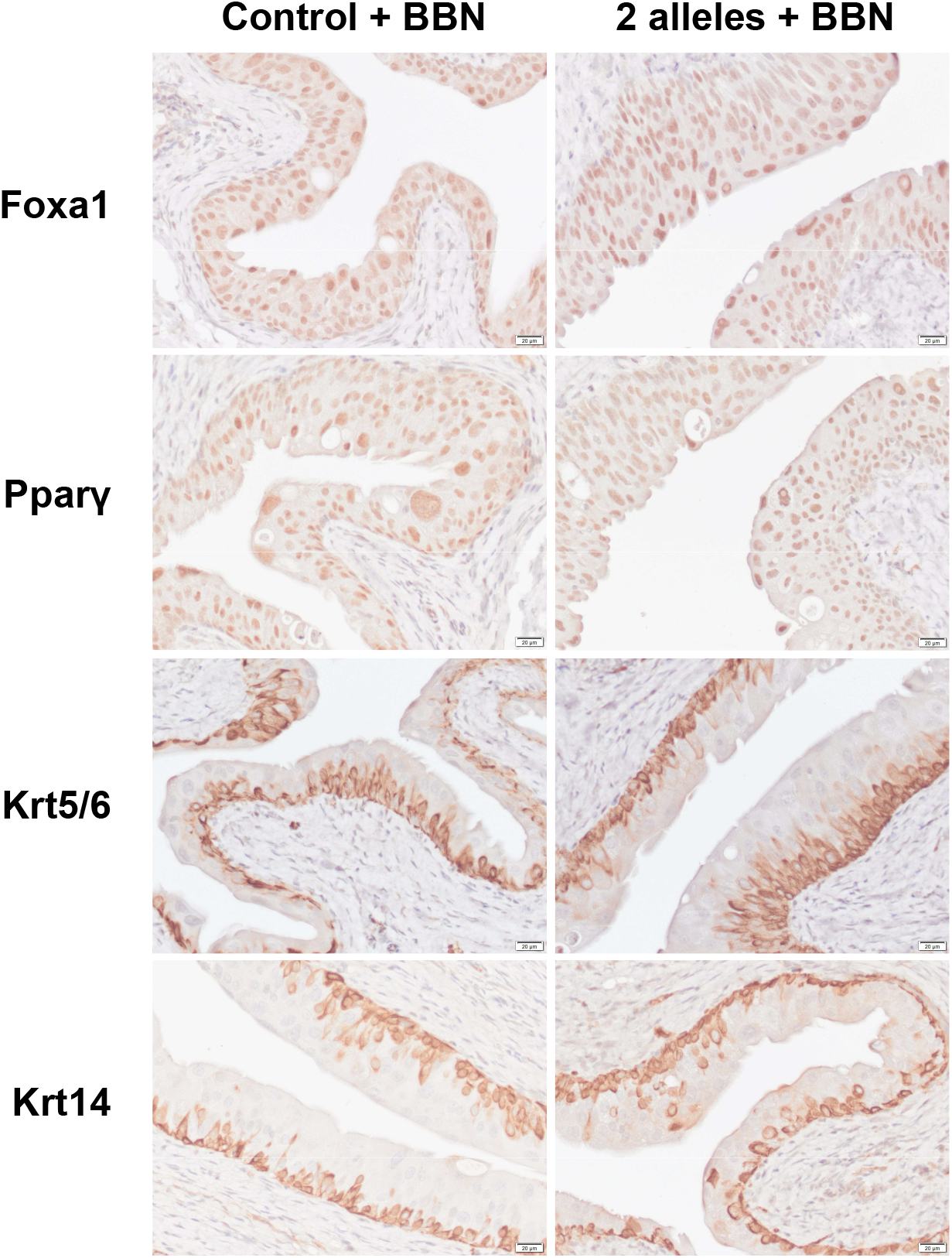
Mutant *Pik3ca* mice maintain similar expression of luminal and basal markers compared to control mice following exposure to BBN. Immunohistochemical staining of luminal (Foxa1 and Pparγ) and basal (Krt5/6 and Krt14) differentiation markers. Mice carrying two alleles of mutant *Pik3ca* exhibit similar expression patterns compared to control mice with high Foxa1 and Pparγ expression, and Krt5/6 and Krt14 expression limited to the basal cells. Scale bar = 20 µm.

### *PIK3CA* mutation is not associated with increased pathway activation in invasive human bladder cancer

The impact of *PIK3CA* mutation in human BC was investigated using TCGA BC cohort primarily comprised of invasive tumor samples.^22^ A lollipop plot was constructed and illustrates E542K, E545K, and H1047R are the three most common mutations (E542K n = 18, E545K n = 28, and H1047R n = 6) (Figure 6A). ssGSEA analysis was then performed to determine the impact of mutant *PIK3CA* on PI3K/AKT/MTOR pathway activation. Tumors with mutant *PIK3CA*, tumors with wildtype *PIK3CA*, and normal tissue samples were compared (p-value = 0.0158; Kruskal–Wallis) (Figure 6B). Follow-up Wilcoxon rank sum pairwise tests showed the scores of tumors with wildtype *PIK3CA* were similar to those of normal tissue (Normal vs. *PIK3CA* WT p = 0.29). However, the scores of tumors with mutant *PIK3CA* were significantly higher than those of tumors with wildtype *PIK3CA* and normal tissue (*PIK3CA* WT vs. *PIK3CA* MUT p = 0.011 and Normal vs. *PIK3CA* MUT p = 0.030). Additionally, tumors with E542K, E545K, and H1047R mutations were highlighted for visual comparison and appear to have similar pathway activation levels to each other. Further ssGSEA analysis was then conducted on a restricted sample set in which samples that harbored oncogenic hotspot mutations within other genes of the pathway (*FGFR3, ERBB2, ERBB3, HRAS, NRAS, PTEN*, and *AKT1*) were removed (Figure 6C). In this analysis, no significant differences were observed among the groups (p = 0.211; Kruskal–Wallis) and Wilcoxon rank sum p-values were as follows: Normal vs. *PIK3CA* WT p = 0.312, Normal vs. *PIK3CA* MUT p = 0.110, and *PIK3CA* WT vs. *PIK3CA* MUT p = 0.189. Tumors with E542K, E545K, and H1047R mutations were again highlighted for comparison and pathway activation levels remained comparable to each other. Overall, these data suggest that *PIK3CA* mutations do not have significant influence on pathway activation in invasive BC.

**Figure 6.**
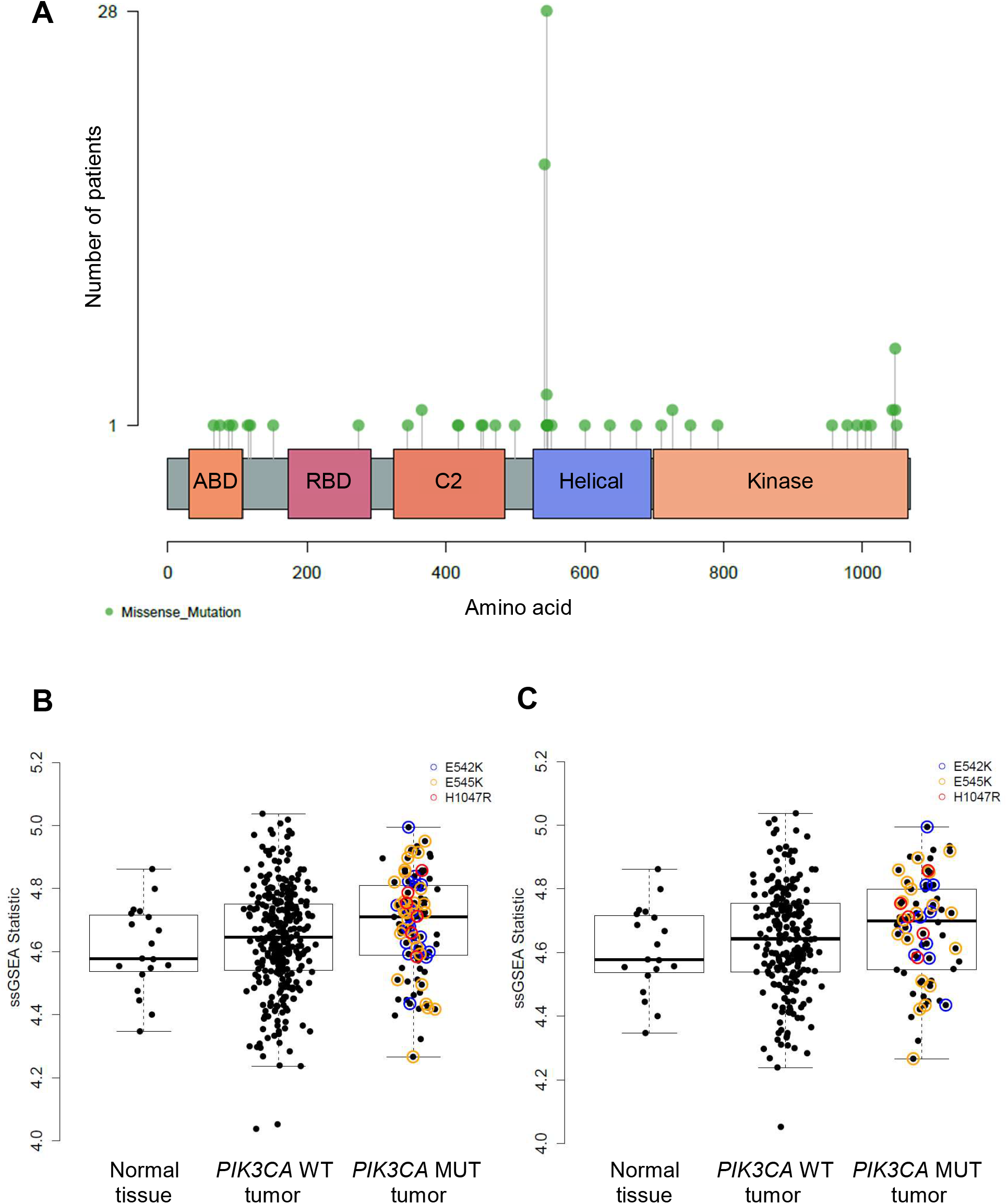
*PIK3CA* mutations in invasive human BC are not associated with increased PI3K/AKT/MTOR pathway activity. (A) Lollipop plot illustrating the number of patients within TCGA BC cohort harboring *PIK3CA* missense mutations and their positions relative to protein domain (E542K n = 18, E545K n = 28, and H1047R n = 6). Abbreviations: ABD – adaptor binding domain, RBD – RAS binding domain. (B) Boxplot illustrating ssGSEA statistic scores of TCGA BC cohort determined by analyzing the PI3K/AKT/MTOR signaling pathway gene set. Normal tissue samples, tumor samples with wildtype *PIK3CA*, and tumor samples with mutant *PIK3CA* were compared (p = 0.0158; Kruskal–Wallis). Pairwise tests using the Wilcoxon rank sum test show the scores of tumors with wildtype *PIK3CA* are similar to those of normal tissue (Normal vs. *PIK3CA* WT p = 0.29). The scores of tumors with mutant *PIK3CA* are significantly higher than those with wildtype *PIK3CA* and normal tissue (*PIK3CA* WT vs. MUT p = 0.011 and Normal vs. *PIK3CA* MUT p = 0.030). Additionally, E542K, E545K, and H1047R amino acid mutations are highlighted for comparison. (C) Boxplot illustrating the same analysis shown in (B), however, samples that also harbored oncogenic hotspot mutations within other genes of the PI3K/AKT/MTOR pathway were removed to focus the analysis on the impact of *PIK3CA* mutations on pathway activation. No significant differences are observed among the three groups (p = 0.211; Kruskal–Wallis). Wilcoxon rank sum p-values are as follows: Normal vs. *PIK3CA* WT p = 0.312, Normal vs. *PIK3CA* MUT p = 0.110, and *PIK3CA* WT vs. MUT p = 0.189). E542K, E545K, and H1047R mutations are again highlighted for comparison.

## Discussion

Activating mutations within the PI3K/AKT pathway are common in many cancers, making pathway components attractive therapeutic targets. More specifically, mutations within *PIK3CA* are frequent in several cancer types including breast cancer and BC.^17, 31^ Mutations in *PIK3CA* appear to be an early event in BC development as they are frequently detected in early stage disease.^16^ The three most common activating *PIK3CA* mutations in BC are E542K, E545K, and H1047R. These mutations occur in the helical (E542K and E545K) and kinase (H1047R) domains and have been found to increase cell motility and migration in normal human urothelial cells.^32^ However, a transgenic mouse model of activating mutations in *Pik3ca* in BC does not currently exist. Coupled with the relative lack of research and availability of models for early stage BC, this is a significant gap in the field. To address this, a *Upk2-Cre/Pik3ca*^*H1047R*^ mouse model was created which is the first model to express mutant *Pik3ca* within the urothelium, thus providing a novel model of BC tumorigenesis.

*Upk2-Cre/Pik3ca*^*H1047R*^ mice were bred and genotyped to identify mice carrying *Upk2-Cre* and one or two alleles of *R26-Pik3ca*^*H1047R*^. To verify the PI3K/Akt pathway was being activated in these mice, IHC was conducted for p-Akt at residues T308 and S473 (Figure 1). Following PI3K activation, Akt is partially activated by phosphorylation at residue T308, then an additional phosphorylation at residue S473 allows for full activation of Akt, making these residues appropriate markers of pathway activation. Results showed nuclear expression of S473 was significantly increased in the mutant mice at 6 and 12 months of age, indicating pathway activation by *Pik3ca*^*H1047R*^. While it was expected that increased cytoplasmic T308 and/or S473 expression would also be observed, the absence of this may have been influenced by scoring or sampling bias. It was noticed that p-Akt appeared to be more prevalent in the urothelium at the neck of the bladder compared to the dome which may have introduced variability contributing to sampling bias. While nuclear expression of T308 was not observed, this was not surprising as it is the first residue phosphorylated and only induces partial activation of the pathway. Overall, with the clear increase in nuclear expression of S473, these results verified the functionality of mutant *Pik3ca* in these mice.

Histological analysis was then conducted to characterize this model. At 6 and 12 months of age, it was observed that mutant mice developed progressive hyperplasia. The thickness of the urothelium was quantified and found to be significantly increased compared to controls (Figure 2). Nuclear atypia was also clearly increased in mutant mice compared to controls and was concentrated within the superficial and intermediate cell populations as was similarly observed with p-Akt staining. As these are the cell populations in which the *Upk2* promoter is active, it was anticipated that the expression of *Pik3ca*^*H1047R*^ would have the greatest effect in these cell populations. This specificity of the *Upk2-Cre* system is one major strength of this study. To increase the number of urothelial cell layers targeted by *Pik3ca*^*H1047R*^, an alternative approach would be to use intravesical delivery of tamoxifen to activate inducible *Cre* systems such as *UBC-Cre*^*ERT2*^ and *Shh-Cre*^*ERT2*^.^33, 34^ This approach would limit the impact of activating *Pik3ca* mutations outside of the bladder while enabling future studies to determine the impact of mutant *Pik3ca* expression within a greater number of cell populations. Future studies could also include prolonged aging of these mice to determine if *Pik3ca* mutation leads to the development of non-invasive BC.

These morphological changes seen in the mutant mice suggest a role for activating *Pik3ca* mutations in urothelial tumorigenesis and support the use of PI3K inhibitors as a targeted treatment option for a subset of non-invasive BC patients. Treatment with PI3K inhibitors is promising and currently used for cancers such as breast and lung cancers.^35^ In fact, studies suggest patients with *PIK3CA* mutations have better response rates to PI3K inhibitors compared to those without *PIK3CA* mutations in breast cancer.^36, 37^ Studies also suggest specific *PIK3CA* mutations may lead to differential responses to PI3K inhibitors; in particular, it has been shown that patients with H1047R mutations had better response rates compared to patients with other or no mutations in *PIK3CA* in various cancers.^38, 39^ Future studies could include use of this model to determine the treatment efficacy of PI3K inhibitors in non-invasive BC with *Pik3ca* mutations. This model could also be used for studying downstream signaling events as well as therapies targeted towards other components of the pathway such as AKT and mTOR.

As BC can be classified into luminal and basal molecular subtypes, we utilized a panel of luminal (Foxa1 and Pparγ) and basal (Krt5/6 and Krt14) urothelial differentiation markers to characterize the *Upk2-Cre/Pik3ca*^*H1047R*^ mice. These mice maintained high expression of Foxa1 and Pparγ and relatively low expression of Krt5/6 and Krt14 (Figure 3), indicating a model of luminal BC tumorigenesis consistent with expression seen in most human non-invasive BCs.

To also assess the contribution of *Pik3ca*^*H1047R*^ to carcinogen susceptibility, we compared control and mutant mice with two *Pik3ca*^*H1047R*^ alleles after 10 weeks of BBN exposure to mice which were not exposed to BBN. Surprisingly, a significant difference in urothelial thickness or nuclear atypia was not observed (Figure 4). This could in part be due to variability induced by BBN. Additionally, no differences in IHC staining for Foxa1, Pparγ, Krt5/6, and Krt14 were observed (Figure 5). Future studies could include the use of different concentrations or exposure times to BBN to elucidate a possible impact of *Pik3ca* mutation on susceptibility to carcinogen.

As *PIK3CA* mutations are seen across all stages of BC, we additionally investigated the impact of these mutations in invasive human BC using a publicly available TCGA BC cohort (Figure 6).^22^ A lollipop plot illustrates E542K, E545K, and H1047R are the three most common mutations as expected. Initial ssGSEA analysis of this cohort suggests patient tumors with mutant *PIK3CA* have significantly higher pathway activity compared to tumors with wildtype *PIK3CA* (p = 0.011; Wilcoxon rank sum test). In an effort to reduce the influence of other mutations on the analysis and therefore focus on *PIK3CA* mutation, further analysis was conducted using a restricted sample set by removing samples with oncogenic hotspot mutations within other genes of the pathway. This analysis suggests, however, that tumors with mutant *PIK3CA* do not have significantly increased pathway activity compared to tumors with wildtype *PIK3CA* (*PIK3CA* WT vs. MUT p = 0.189; Wilcoxon rank sum test). In fact, significant differences were not observed among the groups overall, suggesting pathway activity is not significantly different in tumors in general compared to normal tissue (p = 0.211; Kruskal–Wallis). However, while statistical significance was not reached, median scores are slightly increased in tumors, particularly those with mutant *PIK3CA*. The lack of statistical significance is likely affected by the large variation in scores within the groups which could be due to molecular heterogeneity common in BC. Overall, this data taken together with our experimental mouse model suggests *PIK3CA* mutations may be initiators of pathway activation and tumorigenesis, but become less of a driver of tumor development as additional mutations accumulate and the tumor progresses. Furthermore, it was observed that tumors with E542K, E545K, and H1047R mutations appear to have similar pathway activation levels, supporting the relevance of H1047R in comparison to the other mutations as well as its use in this and future studies.

In conclusion, these data describe *Upk2-Cre/Pik3ca*^*H1047R*^ mice as a novel *in vivo* model of luminal BC tumorigenesis. As *PIK3CA* mutations are commonly detected in early stage BC, this model can be useful in understanding the impact of PI3K/AKT pathway activation in BC tumorigenesis as well as studying the use of targeted therapies including PI3K inhibitors in non-invasive BC treatment.

## Acknowledgments

The authors wish to acknowledge support from the Ruth Heisey Cagnoli Endowment in Urology at Penn State College of Medicine and the Bladder Cancer Support Group at Penn State Health. The authors also wish to acknowledge Jay D. Raman (Department of Urology, Pennsylvania State University College of Medicine) and Amyn M. Rojiani (Department of Pathology and Laboratory Medicine, Pennsylvania State University College of Medicine) for the continued support of our research endeavors.

